# An Algorithm for Sequence Location Approximation using Nuclear Families (ASLAN) Validates Regions of the Telomere-to-Telomere Assembly and Identifies New Hotspots for Genetic Diversity

**DOI:** 10.1101/2022.08.02.502486

**Authors:** Brianna Chrisman, Chloe He, Jae-Yoon Jung, Nate Stockham, Kelley Paskov, Peter Washington, Dennis P. Wall

## Abstract

Although it is heavily relied on to study genetic contributors to health and disease, the current human reference genome (GRCh38) is incomplete in two major ways: firstly, it is missing large sections of heterochromatic sequence, and secondly, as a singular, linear reference genome it does not represent the full spectrum of genetic diversity that exists in the human species. In order to better understand and characterize gaps in GRCh38 and genetic diversity, we developed a method - ASLAN, an Algorithm for Sequence Location Approximation using Nuclear families - that identifies the region of origin of short reads that do not align to the GRCh38. Using unmapped reads and variant calls from whole genome sequencing (WGS) data from nuclear families, ASLAN relies on a maximum likelihood model to identify the most likely region of the genome that a subsequence belongs to, given the phasing information of family and the distribution of the subsequence in the unmapped reads. Validating ASLAN on a synthetically generated dataset, and on true reads originating from the alternative haplotypes in the decoy genome, we show that ASLAN can localize more than 90% of 100-basepair sequences with above 92% accuracy and around 1 megabase of resolution. We then run ASLAN on 100-mers from unmapped reads from WGS from over 700 families, and compare ASLAN localizations to alignment of the 100-mers to the T2T-CHM13 assembly, recently released by the Telomere-to-telomere (T2T) consortia. We find that many unmapped reads in GRCh38 originate from telomeres and centromeres that are gaps in the GRCh38 reference. We also confirm that ASLAN localizations are in high concordance with T2T-CHM13 alignments, except in the centromeres of the acrocentric chromosomes. Comparing ASLAN localizations and T2T-CHM13 alignments, we identify sequences missing from T2T-CHM13 or sequences with high divergence from their aligned region in T2T-CHM13, thus highlighting new hotspots for genetic diversity.

## 2 Background

The human reference genome has been one of the major successes of modern genomics, and has been heavily relied on to study how genetic variation contributes to disease. However, despite its achievements, the current release of the human reference genome (GRCh38) is incomplete in two major ways: it is still missing more than 200Mb (200 million bases) of heterochromatic sequence[3], and it does not represent the full spectrum of genetic diversity that exists in the human species. Because of this incompleteness, reads originating from heterochromatic regions of the genome, or from alternative haplotypes not well represented on the reference, are not analyzed in most genomics studies. On a population-scale, we therefore understand much less about the role these regions play in health and disease, and the amount of genetic diversity present in these regions [16]. Reads originating from these regions - as well as from viruses and bacteria in hosts or NGS reagents [6, 5] - collectively make up the unmapped read space.

We sought to use unmapped reads from whole genome sequences to identify human genomic sequences that do not align to the current human reference genome (GRCh38), and to identify which region of the genome they belonged to, the process by which we refer to as *localization*. Originally, we aimed to develop an algorithm to localize unmapped reads to coarse regions of the genome, with the hope that these regions could function as “bins” and that from these bins, we could ultimately assemble longer contigs that represented alternative haplotypes or sections of the genome missing from GRCh38. We presented a proof-of-concept version of such a localization algorithm for non-repetitive sequences in autosomes, with discussion about a final *de novo* step would look like [7].

While were were developing our algorithm, the Telomere-to-Telomere consortia was finalizing the release of T2T-CHM13, the first gapless assembly of a human genome [21, 18]. Rather than attempting to *de novo* assemble low reliability contigs from scratch, we used our results to validate and analyze against T2T-CHM13, which is likely slated to become the next official human reference genome. The T2T-CHM13 assembly relies on high fidelity long read sequencing (HiFi Sequencing), and novel graph-based algorithms for stitching reads together[21]. Our localizations rely on short-read sequencing and a novel maximum likelihood algorithm that uses sequences from families. Thus, our algorithm for localizing sequences is an orthogonal approach to the T2T’s assembly strategy and presents an excellent opportunity for validating the T2T’s work. We also use our results to understand genetic diversity in relation to the T2T-CHM13 reference genome, especially in regions that were previously gaps in GRCh38 and have thus been understudied. In this paper, we present a modified version of our proof-of-concept algorithm, demonstrate accuracy and precision using a validation dataset, localize subsequences from unmapped reads, and compare the localization results with the results of T2T-CHM13 alignment.

Specifically, we modified our original algorithm to be able to perform on tandem repeats, which are highly present in telomeres and centromeres, and to handle sequences that may originate from the X and Y chromosomes. We present an Algorithm for Sequence Location Approximation using Nuclear families (ASLAN). Named for C.S. Lewis’s lion, ASLAN uses the phasing information of siblings (Peter, Susan, Edmund, and Lucy would have been great additions to our dataset!), and the distribution of a given *k*-mer across the reads of family members in order to build a maximum likelihood model and identify the most probable region of the genome a *k*-mer originates from. We demonstrate ASLAN’s high precision and accuracy on two validation datasets, of which ASLAN can localize over 90% of *k*-mers, with over 90% accuracy, and to around 1 megabase (Mb) resolution. Then, using WGS reads that did not map to GRCh38, we extract 100 bp subsequences and localize them with ASLAN. We show that many of these reads originate from regions with gaps in GRCh38. We compare our localizations to the alignment to the newly released T2T-CHM13 assembly, validating many regions of T2T-CHM13, and identifying regions with potential high levels of genetic diversity.

## 3 Results

### 3.1 ASLAN localizes subsequences using a maximum likelihood model of k-mer distributions and family phasings

We propose ASLAN, an Algorithm for Sequence Location Approximation using Nuclear families. ASLAN requires a large WGS of nuclear families, ideally with multiple children. We use ASLAN with the iHART dataset [26], a large WGS dataset from families with autistic children (4,501 individuals from 1,010 families), that our group curated originally to study the genetic components of autism. To our knowledge, this is one of the largest familial WGS datasets in the world and offers an unique opportunity for family-based analysis such as ASLAN and others [25, 4, 6]. The iHART collection includes the raw WGS from the individuals, as well as the aligned and variant-called data (VCF format) in reference to GRCh38. A detailed description of the iHART data preprocessing process is given in the methods. ASLAN builds a maximum likelihood model that uses family phasing information to identify the genomic region of origin of subsequences extracted from whole genome sequencing reads from this large dataset of nuclear families.

We first phase the children - identify which copy of their mother’s and father’s chromosomes a child inherited a given region - using a hidden Markov model. In this hidden Markov model, the state space is which parental copies of a chromosome a child has at any given region, transitions are recombination points, and observations are the variant calls. A Viterbi algorithm is used to walk through the hidden Markov model to identify the parental copy each child inherited at given genomic region that best explains the variant calls in the family. Next, for a given 100-basepair subsequence (a length that balances uniqueness with likelihood of appearing within a read, see Methods), we extract the number of times it occurred in each individual’s WGS reads. A detailed description of the phasing pipeline is given in the Methods and in Supplementary Methods S1.

We then build a maximum likelihood model that identifies the most likely genomic region for the 100-mer to originate from, given the distribution of the 100-mer and the phasings within and across families, assuming Mendelian inheritance patterns. We validate the accuracy of ASLAN using a synthetic dataset as well as a dataset of real reads from already known locations, and then apply ASLAN to 100-mers extracted from the unmapped read space of the iHART dataset. We illustrate the logic behind ASLAN in Fig. 1 and a full mathematical description in the Methods ad Supplementary Methods S3.

**Figure 1:**
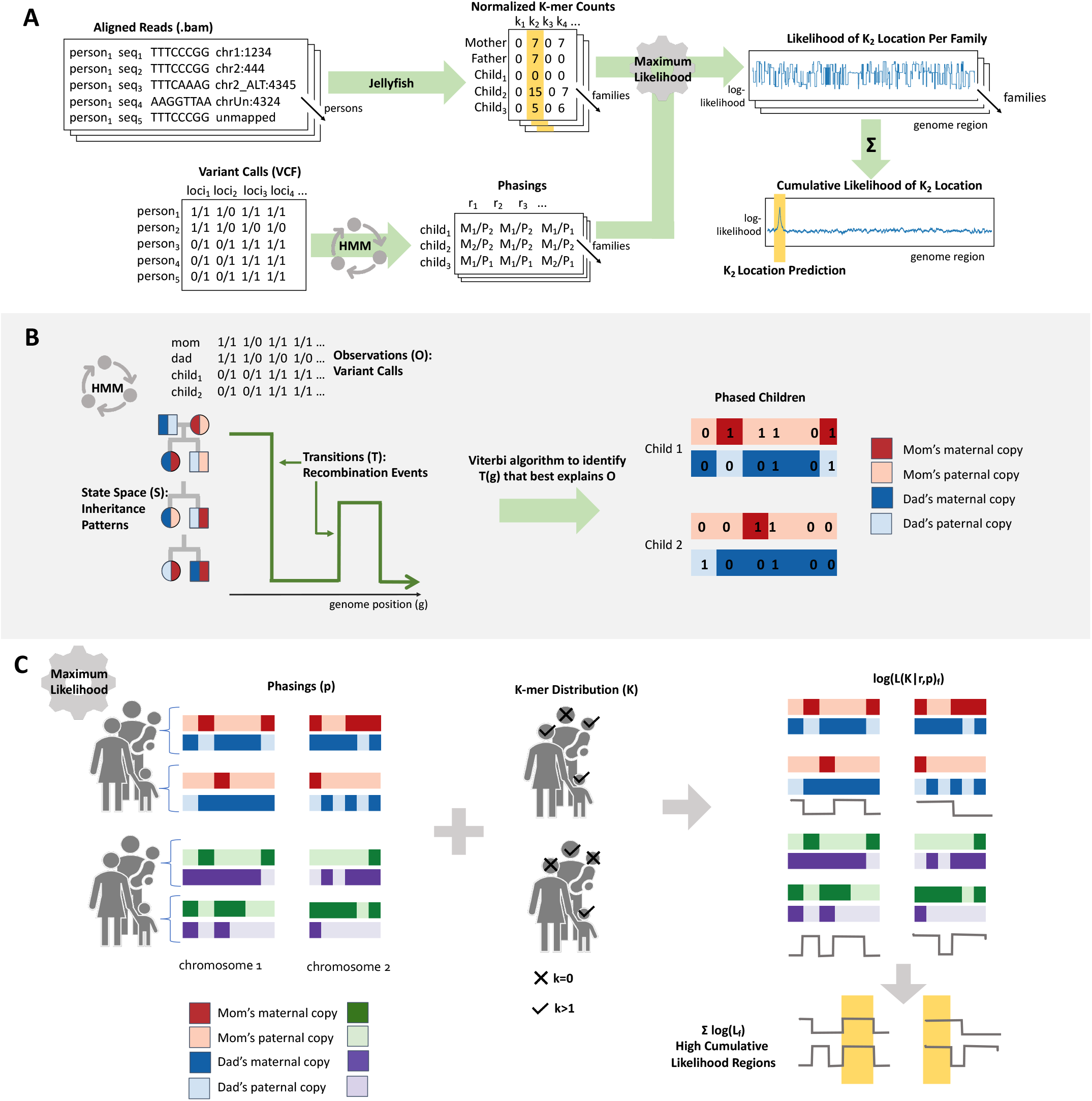
Pipeline for ASLAN and its components. (A) Overall pipeline for extracting *k*-mers, phasing families, and localizing *k*-mers based on phasings and *k*-mer distributions. (B) Simplified schematic of the hidden Markov model used for the phasing algorithm, where the goal is to identify the inheritance patterns and recombination points that best explain the variant calls in a family. (C) Simplified schematic of the the maximum likelihood model which identifies the most likely region of a genome for a *k*-mer to originate from, given the distributions of *k*-mer and phasing patterns within and across families.

### 3.2 ASLAN accurately localizes validation sequences to high precision

We validated ASLAN on two datasets: (1) a synthetically generated dataset and (2) a dataset using *k*-mers extracted from ALT sequences in the decoy genome. Each dataset contained *k*-mer counts and a location label for each *k*-mer. The goals of analyzing ASLAN’s performance on these validation datasets were to understand ASLAN’s performance and to tune ASLAN’s hyperparameters to optimize the balance between precision and accuracy. In this case, precision refers to the size of the region to which ASLAN localizes a 100-mer. For the most part, ASLAN uses biologically computed hyperparameters: the expected number of occurrences of a *k*-mer is computed using the length of the human genome and a sample’s read depth, and inheritance probabilities are derived from simple Mendelian inheritance rules. As described in our sister paper (ref: GENOME/2022/277172), the phasing algorithm uses parameters measured or derived by the literature. The only hyperparameter we use is in the final step of the maximum likelihood model: after we build a cumulative likelihood distribution for the likelihood that a *k*-mer originates from each region of the genome (225,000 possible regions based on the recombination points found within the families), rather than select a single genomic region, we select a range of neighboring regions. The size of this range is governed by the distribution of cumulative likelihoods, and a hyperparameter *λ* (See Fig. 1 and Methods). During validation, we also sought to identify which value of *λ* gave ASLAN the best balance of precision and accuracy.

We first validated our method on the synthetically generated dataset, which simulated theoretical observed counts of *k*-mers given various genomic regions of origin, number of tandem repeats, and prevalences (see Methods). We found that across all tandem repeat numbers, and maximum likelihood hyperparameter values, our algorithm’s performance began to degrade after the prevalence of a *k*-mer became higher than 80%. ASLAN also had poor performance for *k*-mers with prevalences less than 1% (Fig. S2A and C). Smaller values of *λ* resulted in faster degradation of performance in terms of localization ability and accuracy, though smaller values of *λ* also tightened the region length to which *k*-mers were localized. For the most part, ASLAN did not show a bias towards any particular chromosomes, except for chromosome 6 (Fig S2D. This was expected as chromosome 6 contained the most recombination points, and thus had more possible regions available for the algorithm to incorrectly localize a *k*-mer. As shown in Fig. S2, a value of *λ*=.1 gives a good balance of localization ability, accuracy, and precision: our algorithm localizes over 90% of our *k*-mers, is 92% accurate, and has a median localized region length of less than 1Mb.

We then validated our dataset using *k*-mers extracted from alternative sequences in the decoy genome, contigs that are not on the primary reference genome but have been seen in enough individuals to warrant inclusion as alternative sequences in GRCh38. ASLAN performed similarly on this decoy dataset. Changing the *λ* values altered performance similarly to the synthetic dataset. Again, a value of *λ*=.1 provided good balance of localization ability, accuracy, and precision. Using *λ*=.1, ASLAN localized 94% of *k*-mers with 92% accuracy, and a median region length of 1.3Mb. ASLAN again showed a slight preference toward localizing to chromosome 6, which has a disproportionate number of recombination points. There seemed to be some specific hotspots for incorrect localizations, as shown by the spikes in chromosome 6 and 8 in Fig. S3I, which correspond to *k*-mers from a handful of alternative haplotypes being consistently localized to those regions.

Only 24 out of 198 alternative haplotypes had their *k*-mers localized with *<* 90% accuracy. These alternative haplotypes include chr8 KI270813v1 alt, chr12 GL877875v1 alt, chr2 KI270894v1 alt, chr17 KI270907v1 alt, and chr22 KI270878v1 alt.

Interestingly, for *k*-mers from many of these these ‘poorly’ localized haplotypes, our algorithm consistently localized the *k*-mers to another region of the genome. For example, *k*-mers from chr12 GL877875v1 alt consistently were localized to the beginning of chromosome 6, and *k*-mers from chr8 KI270813v1 alt were consistently localized to a region 2Mb downstream of their annotated location. Given these patterns, we wonder if perhaps there are alternative haplotypes located elsewhere in the genome that are homologous to those in the decoy sequence and our algorithm is detecting such.

### 3.3 Many unmapped reads localize to gaps in GRCh38

After validating our algorithm using the synthetic and decoy datasets, we ran ASLAN on 100-mers extracted from the unmapped and poorly aligned reads of WGS 727 nuclear families (we excluded families with half-sibs, twins, or missing a parent). We outline the criteria for what was considered unmapped or poorly aligned in the Methods. We sampled approximately 100 million 100-mers that were found in at least 2 samples, and localized them with ASLAN.

Tuned to the *λ* parameter selected based on the validation data, ASLAN was able to localize 79% of 100-mers with prevalences between 20% and 80% prevalence extracted from the unmapped reads (Fig. 2B-D). For all *k*-mers, which included very low or high prevalence *k*-mers, ASLAN could localize *<*20%. This was expected given that very low or very high prevalences of *k*-mers will not produce enough siblings discordant for a given *k*-mer, which ASLAN depends on for the likelihood model. Many *k*-mers from our model had very low prevalences (see Fig. 2A). We suspect such *k*-mers originate from private familial sequences, sequencing artifacts, and contaminants. Similar to the validation data results, ASLAN localized the unmapped 100-mers to a medium region length of 762,052 bases (Fig. 2E). Since validation showed more reliable performance on *k*-mers between 20% and 80% prevalence, and because ASLAN struggled to localize very low and very high prevalence *k*-mers from the unmapped reads, we focus on these medium prevalence *k*-mers. The rest of our findings only use *k*-mers between 20% and 80% prevalence.

**Figure 2:**
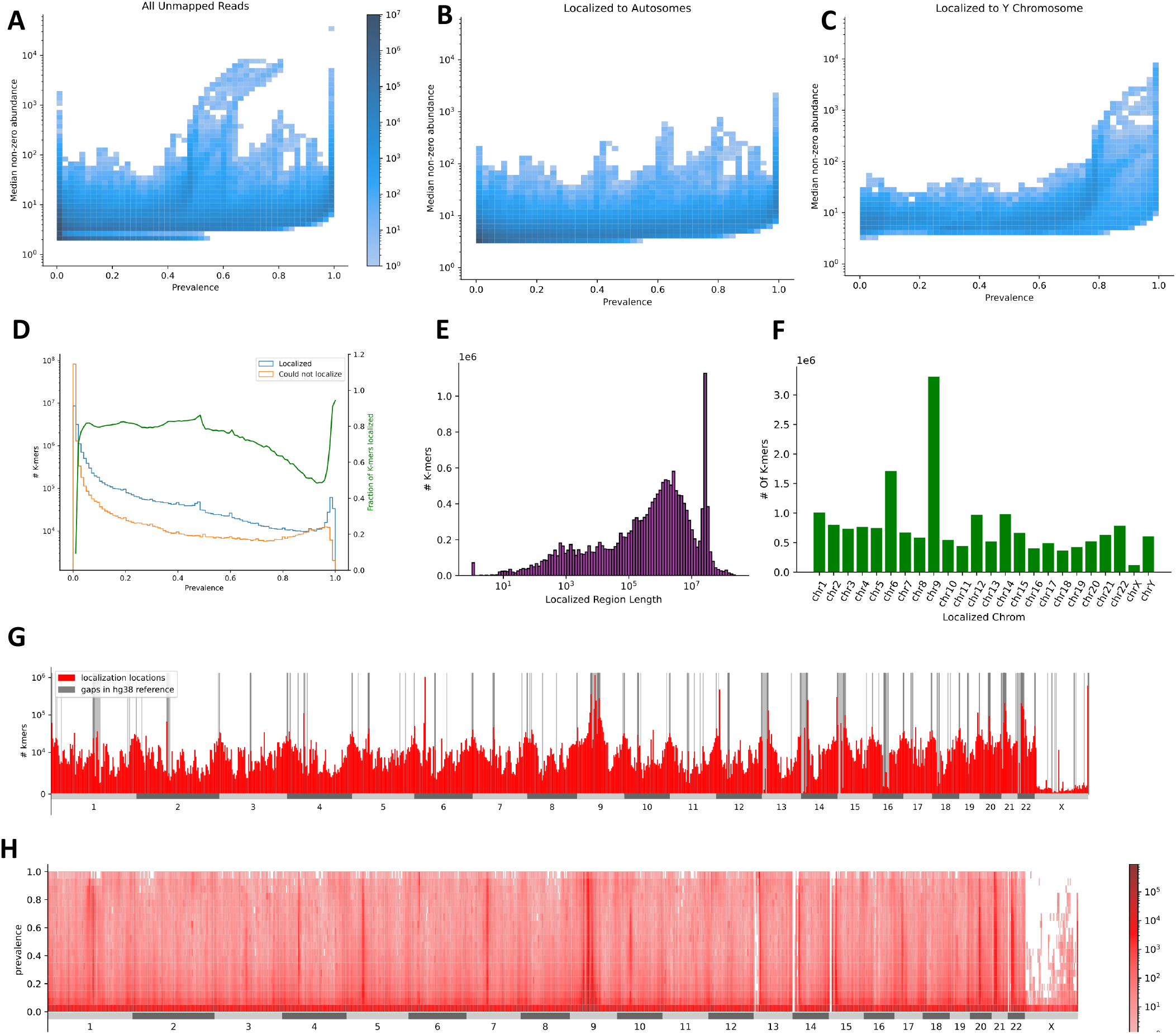
ASLAN performance on unmapped reads. (A) Distribution of prevalence and abundance (median of non-zero counts) for all 100-mers extracted from unmapped reads. (B) Distribution of prevalence and abundance for 100-mers that localized to autosomes. (C) Distribution of male prevalence and abundance for 100-mers that localized the Y chromosome. (D) Number of localized and unlocalized 100-mers, and corresponding localization fraction at various prevalences. (E) Distribution of localized region length. (F) Number of *k*-mers localized to each chromosome. (G) Distribution of localization location in reference to GRCh38, with gaps annotated. (H) Distribution of localization location vs. prevalence.

We found that many unmapped 100-mers localized to regions that included gaps in GRCh38 (Fig. 2G-H), more than expected by chance, as demonstrated using a Monte Carlo simulation shuffling the gap locations to generate a null distribution. We found that 56% of localization regions contained at least one gap present in GRCh38 (p*<* .05) and 49% of localizations had a center point that fell within a GRCh38 gap (p*<*.0001). In particular, we see many *k*-mers localizing to the large gaps representing the centromeres of chromosome 1 and 9. We also see many sequences localizing to gaps that represent the short arms of the acrocentric chromosomes (chromosomes 13, 14, 15, 21, and 22). Even with the highly repetitive nature of heterochromatin, it seems that ASLAN is able to identify many subsequences coming from the heterochromatic gaps in GRCh38. These successfully localized subsequences probably include tandem repeat sequences that are unique to a single region of heterochromatin, unique sequences in flanking regions of repetitive sequences, or repeats in heterochromatin that have developed a variant, giving some frequency of the human population a variant on a repeat.

There are several hotspots where sequences localize that are not gaps in GRCh38 - chr2:89Mb, chr2:175Mb, chr3:167Mb, chr3:175Mb, chr6:33Mb,chr8:13Mb, chr8:18Mb, chr8:136Mb, chr11:1Mb, chr11:127Mb, chr12:11Mb, chr15:20Mb, chr19:55Mb, and chr20:29Mb. We describe the functional relevance of these regions in the Discussion.

### 3.4 Metacentric chromosomes show high concordance between ASLAN localizations and T2T-CHM13 alignment

During the course of our study, the Telomere-to-Telomere consortia released the sequence of the T2T-CHM13 genome, the first fully assembled human genome. While it would be time-intensive and potentially error-prone to rerun the entire iHART and ASLAN pipeline using the far less studied T2T-CHM13 genome as the primary reference, we were well aware of the relevance of T2T-CHM13 to the goals of ASLAN. We decided to align the sequences of the ASLAN-localized 100-mers to the T2T-CHM13 assembly in order to identify regions where ASLAN and the T2T-CHM13 alignment were in high or low concordance, and to identify potential sources of genetic diversity, especially in the gapped regions of GRCh38.

We first analyzed ASLAN and T2T-CHM13 concordance. Our thinking was that for *k*-mers where ASLAN localizations and T2T-CHM13 mappings agree, we can be more certain that these *k*-mers truly originate from the specified region. If T2T-CHM13 alignment score is low, this is probably indicative of some genetic variants between the set of iHART samples and the T2T-CHM13 genome. If ASLAN localizations and T2T-CHM13 mappings disagree, it might be indicative of homology between a T2T-CHM13 region and a different region in a subset of the iHART samples. T2T-CHM13 represents only a single genome, and is important to understand the limits of the single genome, especially in the previously gapped regions where no large scale analyses of structural variation exist.

We found that ASLAN and T2T-CHM13 showed a very high concordance on the metacentric and submetacentric chromosomes (chromosomes 1-12, 16-20, and X). Across the entire genome, 69% of sequences mapped to the same chromosome, using ASLAN and T2T-CHM13 alignment (Fig 3B). For close to 80% of sequences localized to each of the metacentric and submetacentric chromosomes, ASLAN and T2T-CHM13 mappings were concordant: the ASLAN-localized region contained the T2T-CHM13 alignment location (Fig. 3D). Mismatches were most common between the sequences that aligned to the T2T-CHM13 centromeres of chromosome 1 and 9, and the short arms of chromosomes 13-15 and 21-22-the acrocentric chromosomes, which we discuss below. ASLAN localizations of these mismatches were fairly uniformly distributed across the genome (Fig. 3A-B) with no glaring preferences towards certain chromosomes or regions.

**Figure 3:**
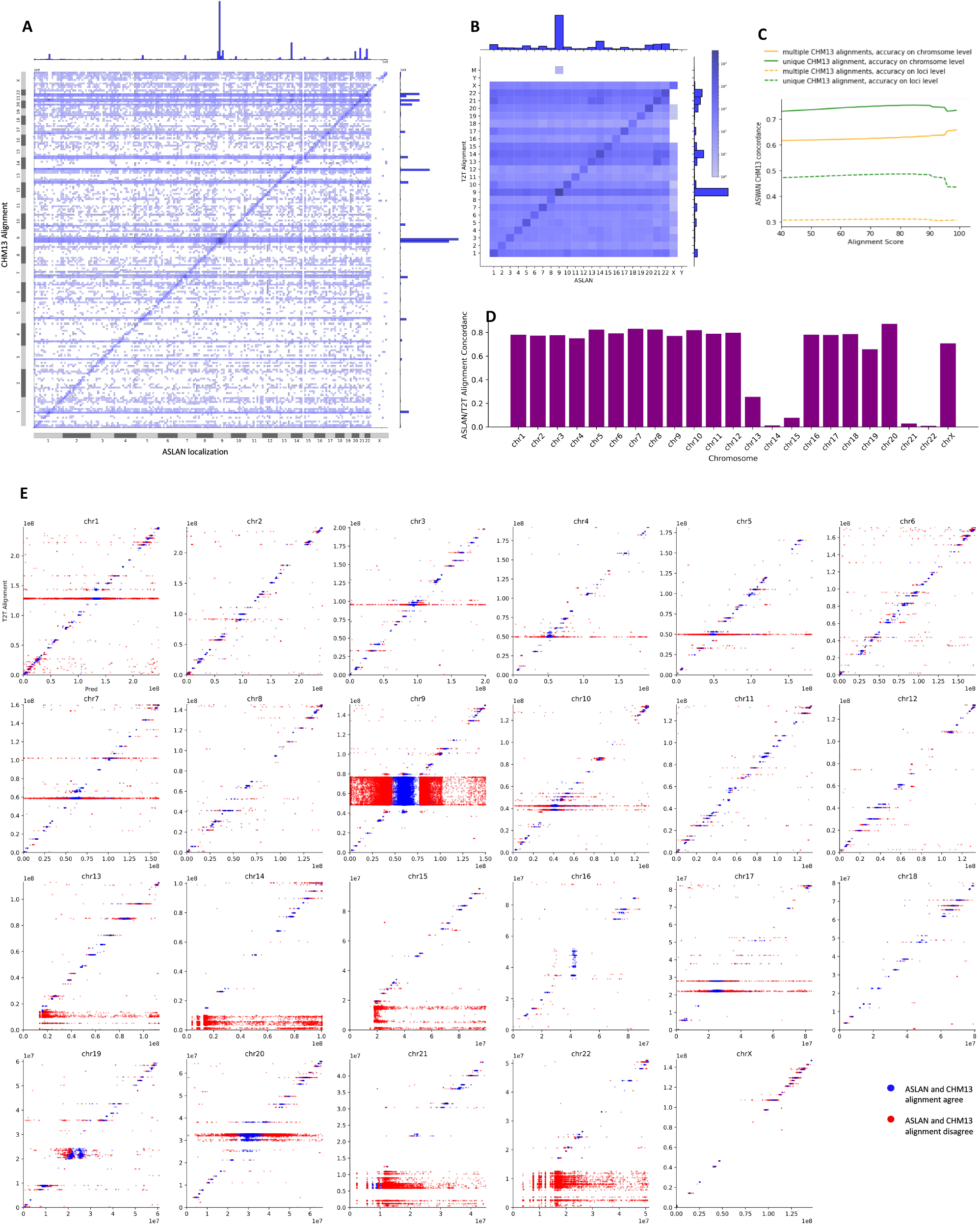
(A) Confusion matrix comparing ASLAN localizations, lifted over to T2T-CHM13 coordinates to T2T-CHM13 alignments of 100-mers extracted from the unmapped reads, binned into 1000 equally sized bins across the genome. (B) Confusion matrix comparing ASLAN localization to T2T-CHM13 alignment, binned by chromosome. (C) Concordance rate between ASLAN localization and T2T-CHM13 alignment, vs alignment score, and colored by whether or not alignment to T2T-CHM13 was unique or not. (D) Concordance rate between ASLAN localization and T2T-CHM13 alignment, vs chromosomes. Acrocentric chromosomes 13-15, and 21-22 show a significantly lower concordance. (E) T2T-CHM13 alignment vs center point of ASLAN localization region, separated by chromosome and colored by colored by whether or not alignment and localization were in concordance.

Disagreement between ASLAN localization and T2T-CHM13 alignment may be caused by one of several reasons. First of all, the ASLAN localization may be incorrect. ASLAN demonstrated 92% on fairly clean data: synthetic data and data from the primarily euchromatic alternate haplotypes in the decoy genome. Even on the validation data, ASLAN still has an 8% error rate and it is possible that ASLAN has slightly degraded performance on the *k*-mers extracted from the unmapped reads, the majority of which seem to originate from heterochromatin. Secondly, it is possible that a *k*-mer originates from a haplotype not be well-represented on T2T-CHM13, but that does appear in a fraction of the iHART genomes. In this case, the T2T-CHM13 alignment may be incorrect, and the ASLAN localization correct. If the *k*-mer is highly divergent from any region in T2T-CHM13, we may see a very low alignment score when mapping to T2T-CHM13. However, if the true region of origin has homology to a different region in T2T-CHM13, it might be impossible to determine whether the ASLAN localization or T2T-CHM13 is correct. It seems possible that some of the *k*-mers in our dataset are being mismapped to regions in T2T-CHM13. Fig. 3C, shows that ASLAN-T2T-CHM13 concordance does seem to go up slightly as alignment score increases. Furthermore, ASLAN vs. T2T-CHM13 concordance is much higher for *k*-mers that uniquely aligned to a single region on T2T-CHM13 compared to those that had multiple similarly-scored alignments. We also see that *k*-mers where ASLAN localizations and T2T-CHM13 alignments disagreed had lower alignment scores (Fig. 4D-E). Concordance between ASLAN localizations and T2T-CHM13 alignment showed a very bimodal distribution across chromosomes (Fig. 3D). While the metacentric and submetacentric chromosomes all showed close to an 80% concordance with the T2T-CHM13 alignments, the acrocentric chromosomes (13-15, 21-22) showed less than a 20% concordance. We also saw a much lower concordance in the centromeric and short arms regions of the metacentric and submetacentric chromosomes (Fig. 3E). These two phenomena are likely related and may be caused by of a few reasons, assuming the T2T-CHM13 assembly is correct, as it was validated using several different sequencing modalities. We discuss possible reasons for disconcodance and the limitations of both ASLAN and T2T-CHM13 further in the Discussion.

**Figure 4:**
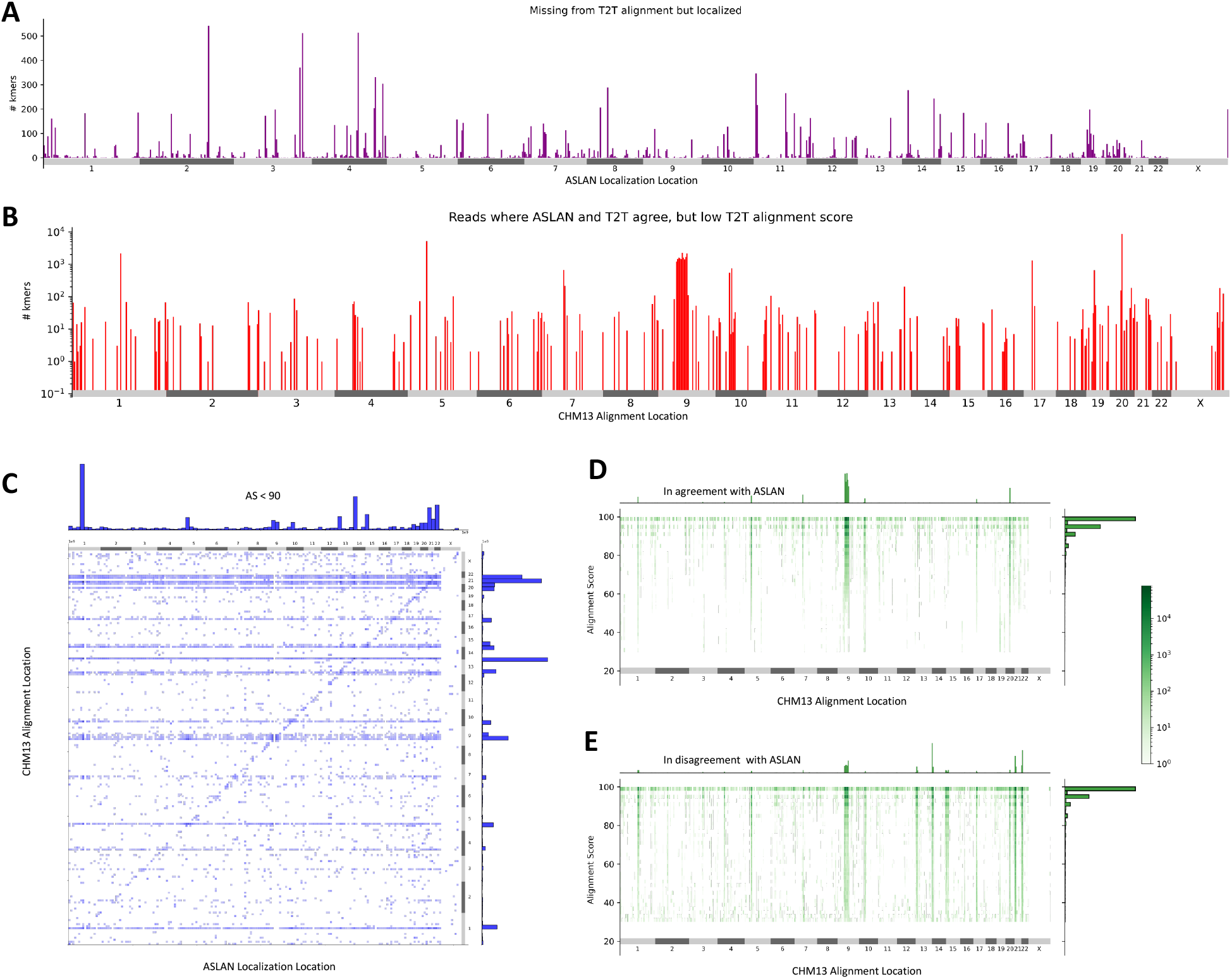
(A) Distribution of reads that failed to align to the T2T-CHM13 assembly, but that were successfully localized via ASLAN. (B) Distribution of reads where the localization region predicted by ASLAN contained the location the read aligned to on T2T-CHM13, but the alignment score was less than 90. These regions may indicate new hotspots for genetic diversity. (C) Joint plot of regions where ASLAN localization and T2T-CHM13 alignments were in disagreement with one another, and the T2T-CHM13 aligmnet score was *<*90 These may indicate sequences that are not represented in the T2T-CHM13, but that are highly homologous to a different region in T2T-CHM13. (D)-(E) Loci and alignment score distribution between the T2T-CHM13 alignments for *k*-mers with ASLAN localizations and T2T-CHM13 alignments (D) in agreement with each other and (E) in disagreement with each other. *k*-mers in disagreement have significantly lower alignment scores, suggesting that imperfect alignments to T2T-CHM13 may actually be originating from a a human genome sequence not well represented on T2T-CHM13.

### 3.5 Comparison of ASLAN localizations and T2T-CHM13 alignments highlights new hotspots for genetic diversity

While T2T-CHM13 is the first assembly of a full human genome, it is still the sequence of a single genome. We next compared ASLAN’s localization results to the T2T-CHM13 alignment in order to analyze genetic variation, particularly in the regions of the genome that have been understudied due to their absence in the reference genome. We looked at three types of genetic variation: (1) 100-mers that failed to align to anywhere in the T2T-CHM13 assembly, but that were successfully localized with ASLAN, indicating regions of the human genome where T2T-CHM13 and the genome of other samples are extremely divergent; (2) 100-mers where T2T-CHM13 alignment and ASLAN localization agree, but that have a low T2T-CHM13 alignment score, indicating regions where large amounts of SNPs or simple genetic diversity exists across samples; and (3) regions where ASLAN localizations and T2T-CHM13 alignments do not agree, that had low T2T-CHM13 alignments scores, suggesting regions where T2T-CHM13 alignment may not be reliable because there are homologous sequences in regions of the human pan-genome not captured on T2T-CHM13.

First of all, we found that a small (1.5%) percentage of our unmapped *k*-mers that localized successfully with ASLAN failed to align to T2T-CHM13 at all (Fig. 4A). Of the total 20%-80% prevalence *k*-mers that did not align to T2T-CHM13, ASLAN was able to localize 88% of them. According to the ASLAN localizations, these *k*-mers seemed to originate from the Y-chromosome (37%) and the autosomes (63%). It makes sense that many of these *k*-mers originate from the Y chromosome: T2T-CHM13 is from a hydatidiform mole, and so the assembly does not contain a Y chromosome, though the T2T is in the works of fully assembling a Y chromosome from the Genome in a Bottle collection. The T2T-CHM13-unaligned *k*-mers that localized to autosomes with ASLAN seem to originate from specific regions of the genome.

These may indicate hotspots for genetic diversity, where a sequence exists in an iHART haplotype that is not well-represented on the singular T2T-CHM13 genome. These hotspots include regions around chr2:176Mb, chr3:171Mb, chr3:177Mb, chr4:117Mb, chr11:7Mb, and overall chromosome 19 (in GRCh38 coordinates). Unlike the hotspots for localized reads in general, these hotspots for localized but unaligned reads do not seem to have any relationship with known genetic diversity, as catalogued by dbVar and the alternative sequences in the decoy. Notably, within the hotspot around chr3:177Mb is the repeat-containing gene TBL1XR1. Mutations in TBL1XR1 have been previously associated with autism spectrum disorders [23]. As the iHART dataset is from multiplex autism families, it is possible that ASLAN is localizing a structural variant in TBL1XR1 that is not represented on T2T-CHM13, but may be more frequent in families with autism.

We also found many regions where ASLAN localization and T2T-CHM13 alignment seemed to be in agreement, but where the T2T-CHM13 alignment had a relatively low alignment score (*<*90) (Fig. 4B). With short query sequences, it can be difficult to tell whether a low alignment score indicates that a query sequence does in fact originate from the region of the genome it aligned to but has variation compared to the reference, or if it originates from another region from the genome entirely, possibly from a genomic region not well represented on the reference genome used in alignment. Given that ASLAN localizations and T2T-CHM13 alignment were the same in these cases, we assumed that these *k*-mers were the former, and truly did belong to the region that they aligned to in T2T-CHM13. The low alignment score would then be caused by genetic variation relative to the T2T-CHM13 reference sequence.

Again, we saw hotspots for this type of genetic variation. Even more dramatic than for the *k*-mers that failed to align to T2T-CHM13, the number of low-AS *k*-mers with ASLAN and T2T-CHM13 concordance differed by over 5 orders of magnitude across different regions of the genome (Fig. 4B). Specifically, we see hotspots within the centromeres of chromosomes 1, 5, 7, 9, 10, 17, and 20, and near loci chr8:137Mb and chr13:96.5Mb. The alignment near chr8:137Mb is within the long non-coding RNA sequence, LINC02055 and the alignment near chr13:96.5Mb is within the gene HS6ST3.

Finally, we analyzed *k*-mers where ASLAN localizations and T2T-CHM13 alignment did not agree, and T2T-CHM13 AS was low. We discussed these types of sequences in the previous section but revisit them briefly. The *k*-mers are a bit trickier to analyze: on one hand, they could just be a sample of the *k*-mers that ASLAN incorrectly (as seen in the validation dataset, ASLAN with the chosen hyperparameter, does localize around 8% of *k*-mers incorrectly. On the other hand, these *k*-mers could be originating from a genomic sequence not well captured in T2T-CHM13, with the low alignment score suggesting that the aligned location is not the true origin of the *k*-mer. As we discussed previously, the lower alignment scores for *k*-mers that were discordant between T2T-CHM13 alignment and ASLAN localization compared indicates that T2T-CHM13 alignment may be incorrect for at least some of these mismatched mappings. Particular hotspots for discordance and low alignment scores are the *k*-mers that aligned to the T2T-CHM13 centromeres in chromosome 1, 5, 9, 10, and 17, and small regions within the short arms of the acrocentric chromosomes (Fig 4C).

As most modern genomicsts understand, alignment a sample’s reads to a singular reference genome, even one as complete as T2T-CHM13, does not necessarily give us a full understanding of sample’s variants: It may be difficult to distinguish between misalignment and divergence from the reference sequence, and some sequences within a sample may be poorly represented on the reference genome. In the genetically diverse hotspots we highlighted, particular care should be taken when performing alignment to T2T-CHM13, and alternate sequences may be needed to ensure adequate representation of genetically diverse regions in the human genome.

## 4 Discussion

Taking advantage of the unique family structure of the iHART dataset, we built an algorithm that uses phasing patterns across families to identify probable regions of origin of unmapped sequences. ASLAN shows high performance on validation datasets, and localized many unmapped that seemingly originate from gaps in GRCh38. Comparing ASLAN localizations with T2T-CHM13 alignments, we identified several regions of interest that may warrant further study in future with high fidelity human genome assemblies: regions where ASLAN localizations and T2T and T2T-CHM13 alignments were in agreement but where the low T2T-CHM13 alignment scores may indicate hotspots for genetic diversity, as well as regions where ASLAN localizations and T2T-CHM13 alignments disagreements may indicate sequences that are homologous to a T2T-CHM13 region, but they themselves not well-represented on the T2T-CHM13. As the field of pan-genomics hones in on developing a human reference genome that encompasses genetic diversity, these genetically diverse regions we mention in our result may be regions of the genome to prioritize.

### 4.1 Localization Hotspots

As described in the Results, ASLAN localizes reads to many localization hotspots. Localization locations are enriched for gaps in GRCh38, and non-gapped locations may genetic diversity hotspots where the single reference genome does not do a good job of capturing all sequences.

Some of these hotspots (chr2:89Mb, chr8:13Mb, chr15:20Mb) contain a fix patch, an known incorrectly assembled region of genome that will be updated in the next release of the reference genome. The subsequences localized to this region probably went originally unmapped due to the incorrect assembly on GRCh38. Note that we did not use the fix patches in our original alignment, as is the standard in alignment pipelines because often fix patches indicate a more complicated change than a simple inserted or alternate sequence, such as a change in coordinate system.

Several of these regions are catalogued hotspots for genetic diversity. The either have alternate haplotypes catalogued in the reference genome (chr 6:33Mb, chr19:54Mb, and chr12:11Mb), contain high frequency structural variations catalogued by dbVar [13] (chr8:136Mb, chr11:1Mb, chr11:127Mb), and are likely home to additional uncatalogued complex variation.

The localization hotspot on chromosome 6 (around 33Mb) is a known hotspot for genetic diversity - it lies within the HLA gene set, the most diverse region of the genome across individuals. The decoy genome lists 6 specific alternative haplotypes in this region, but it is estimated there are many orders of magnitudes more across the human population [8]. These sequences were extracted from reads that did not align well to the primary reference genome nor the decoy contigs; therefore ASLAN is localizing HLA subsequences that are not well represented anywhere on GRCh38. Similarly, chromosome 19 (around 54Mb) contains over 30 alternative haplotypes. These region is home to the killer immunoglobulin receptor (KIR) gene family, involved in also involved in immune function and known to be extremely diverse across the human population [27]. Finally, the hotspot on chromosome 12 (around 11Mb) has two additional alternate haplotypes associated with it, and likely more alternative haplotypes not catalogued in the decoy genome.

The hotspot on chromosome 8 around 136Mb is a known hotspot for structural variation, especially within non-European populations, with numerous insertions and deletions with high allele frequency (*>* 0.2) catalogued on dbVar in the long non-coding RNAs (LINC 2005 and RP11-149P24.1) The hotspots on the start (1Mb) and end (127Mb) of chromosome 11 also show evidence of widespread genetic diversity. Within the first 1Mb of chromosome 11, there are 3 alternative haplotypes in the decoy genome, and likely even more variation not catalogued in the decoy. At the end of chromosome 11 (127Mb), there is widespread high frequency structural variation catalogued in dbVar, particularly in the KIRREL3 gene and the lnRNA LOC101929473.

### 4.2 Discordance Between CHM13 and ASLAN

Although reads localized to the metacentric and submetacentric chromosomes have high degrees of concordance using T2T-CHM13, the acrocentric chromosomes do not perform as well. There are a number of possible explanations for this. First of all, although ASLAN designed and works well for tandem duplications that occur in only a single region of the genome, it is not designed for segmental duplications, especially if they occur in multiple regions of the genome or if there is a recombination point between repeats. This may be the case for SINEs, LINEs, LTRs, and other non-tandem repetitive elements. Secondly, our phasing algorithm relies on variant calls, in relation to the reference genome. We cannot therefore, precisely identify recombination events that happen in gaps in the reference genome. If a recombination event happened within a gap in GRCh38, our algorithm would at best, identify that recombination point to occur right after or right before the gap. If two recombination events occurred within a gap, our phasing algorithm would miss the recombination events entirely. Additionally, PCR amplification bias of certain low GC-content regions in the centromeres [1] may invalidate the assumption that observed subsequence count follows a Poisson distribution uniform across the genome.

Finally, like many sets of reference genomes, T2T-CHM13 and GRCh38 have differences in their coordinate systems: a contiguous segment of genome in GRCh38 may not translate to a contiguous segment of genome in T2T-CHM13. T2T-CHM13 has deletions, insertions, and translocations relative to GRCh38. Thus, a region predicted by ASLAN, a contiguous sections of GRCh38, may include several non-contiguous regions of T2T-CHM13, some of which may be on different chromosomes. We used the software package `liftOver` [10] and the chain files provided by the T2T to translate our GRCh38 localizations in to T2T-CHM13 coordinates to compare to T2T-CHM13 alignments. Lifting over each base contained in a large region usually results in an output that is split across multiple regions, and often even multiple chromosomes, of a genome. To avoid confusion with this, we lifted over only the ends of our region, which can be specified in the `liftOver` arguments and was developed to lift over large regions such as BACs. Therefore, our contiguous regions of GRCh38 were lifted over to contiguous regions of T2T-CHM13, even though it was possible they contained sequences of T2T-CHM13 outside the output regions. If a 100-mer originated from one of these sequences, there may be a mismatch between its T2T-CHM13 alignment and the ASLAN localization when translated to T2T-CHM13 coordinates, even if ASLAN actually performed correctly.

Similar to several other studies [16, 9], our results showed that the telomeres and centromeres in the human genome are major sources of genetic variation. The combination of long-read technology and the T2T-CHM13 reference genome will likely open up new doors for population-scale studies on roles these satellite sequences to human health and disease [17]. Some the other highly diverse regions we found fell into regions where others have observed high levels of sequence divergence or structural variation, as catalogued in dbVar and the alternative haplotypes in the decoy genome. We also identified additional candidate hotspots for genetic variation that to our knowledge have not been previously characterized as major contributors to human genetic diversity, including regions within the TBL1XR1 and HS6ST3 genes and the long non-coding RNA LINC02055.

### 4.3 Prioritizing Studies of Global Genetic Variation

Regrettably, although some of the goals of this study were to characterize genetically diverse regions of human genome, the iHART dataset falls prey to many of the diversity and representation issues that modern genomics has been grappling with. The iHART dataset, which encompasses a subset of samples from Autism Genetic Resource Exchange (AGRE) biobank [12], primarily contains individuals of European descent: 83.5% of our participants self-identified as white, 2.7% as Asian, 2.9% as Black, less than 1% each Alaskan native/American Indian and Native Hawaiian/Pacific Islander, 5% mixed race, 5% mixed race and 4% unknown. Autism research poses a particularly challenging set of circumstances in data ascertainment, where socioeconomic and demographic factors play a major factor in the availability of resources for diagnosis, treatment, and connections to research studies[2]. Nevertheless, the iHART dataset is not only limited when it comes to understanding autism genetics of diverse populations, but using ASLAN on iHART only gives us insight into genetic diversity of a small subset of the global population. While there are several more representative WGS datasets currently in the making, ASLAN’s requirements are fairly niche in that it requires large numbers of multi-children families. To our knowledge, iHART is the one of the only datasets of this kind.

On the bright side, regions of the genome that are genetically diverse within populations also tend to be genetically diverse across populations, as these hotspots may be in regions that are subject to higher de novo mutation rates or reside in genetic elements that can handle higher mutational loads [19, 24, 20] Though there may be additional hotspots for genetic diversity in highly diverse populations such as West Africans, our newly highlighted regions of genetic diversity (Fig. 4A-B) within the relatively ethnically homogeneous iHART population likely generalize to other populations as well. These hotspots of genetic diversity are thus important areas to focus on on as long-read technology makes it possible to quickly assemble partial genomes. If these regions are candidates for disease-association studies, the studies may require larger sample sizes or need to include in-depth analysis of structural variation.

## 5 Methods

### 5.1 Data Pre-processing

#### 5.1.1 iHART Dataset, GRCh38 Alignment, Variant-calling

We used the iHART WGS collection [26], a dataset of multiplex autism families, containing 1,006 families and 4,610 individuals. Individuals were sequenced at 30x coverage using Illumina’s TruSeq Nano library kits, reads were aligned to build GRCh38 of the reference genome and decoy contigs using bwa-mem [14], and variants were called using GATKv3.4. Only biallelic variants that passed GATK’s Variant Quality Score Recalibration (VQSR) were included in analysis.

#### 5.1.2 Extracting and Realigning Unmapped and Poorly Mapped Reads

We excluded secondary alignments, supplementary alignments, and PCR duplicates from downstream analyses. We extracted reads from the iHART genomes that were unmapped or mapped with low confidence. Low-confidence reads were defined as reads marked as improperly paired and with an alignment score below 100. We used alignment score rather than mapping quality in order to select for reads were likely not true alignments to the human reference genome, rather than for reads that had ambiguous alignments to GRCh38.

We used Kraken2 [28] to align the unmapped and poorly aligned reads to a the Kraken default (RefSeq) databases of archaeal, bacterial, human (GRCh38.p13), and viral sequences [22]. These references databases were accessed on Feb 16, 2021. Kraken2 was run on the unmapped and poorly mapped reads from each sample, using the default parameters. We then removed the reads that Kraken was able to classify as archaeal, bacterial, viral, or human reference sequences to leave us with the ultimately unmapped reads. This corresponded to a median of 4.4 million reads per sample, or 0.6% of the total number of reads.

### 5.2 Localization Algorithm

#### 5.2.1 Phasing

Phasing refers to the use of an individual’s genetic data to determine which sequences or variants in their genome were inherited from their mother and which from their father. During meiosis, each parents’ two copies of each chromosome are combined in large blocks to form a new chromosomal copy which is then inherited by the child. In phasing, the goal is to identify which parental copy was inherited by each child at every position of the genome. We used a hidden Markov model (HMM) developed in our related work to phase the families in the iHART dataset. The phasing algorithm ultimately outputs for each region of the genome, whether a child inherited copy 1 or copy 2 of their mother’s genome in the form of (*m*_1|2_, *p*_1|2_). We describe the mechanics of the phasing algorithm fully in our other work (ref: GENOME/2022/277172), and briefly in the Supplementary Methods (S1).

#### 5.2.2 Extracting K-mer Counts

Although the read lengths in the iHART dataset were 150 bp, attempting to localize an entire sequence of a read runs the risk that a sample’s genome contains a specific 150 bp sequence, but no reads from that sample originate from that exact region. Therefore, we instead extracted subsequences from the raw reads. We aimed to extract a subsequence that was short enough to nearly guarantee that if a sample’s genome contained the subsequence, the sample’s reads would as well, but not so short that a subsequence might occur many times across various regions of a genome. To accomplish this, we chose a k-mer length of 100bp. The full derivation is shown in the Supplementary Methods (S2) and Fig. S1.

From the reads of interest (either reads mapping to the decoy genome in the validation pipeline, or poorly aligned and unmapped reads), we converted these alignments back to raw reads using `bam2fastq` then the fast multi-threaded *k*-mer counter `jellyfish` [15] to extract and count subsequences of 100bases from the reads. To reduce the amount of low abundance contaminants and reads originating from sequencing errors,for each sample, we extracted and counted only non-singleton *k*-mers. Again, using `jellyfish` we catalogued and counted the 100mers that appeared at least twice in at least two samples in the iHART dataset.

#### 5.2.3 Maximum Likelihood Model

We previous developed and validated a proof-of-concept algorithm to localize 100-bp *k*-mers extracted from 150 bp reads [7]. We review the mathematics of this maximum likelihood model, and discuss the modifications that we added in order to allow localization of tandem repeats and for sequence originating from the sex chromosomes.

The goal of the maximum likelihood model is as follows: For each *k*-mer, we wish to find it’s corresponding location (region *r*) in the genome that best explain the distribution of the *k*-mer counts in all of the families. We define the distribution of a given *k*-mer in all samples as **K**, and the distribution of a given *k*-mer in family *f* as **K**_**f**_. Therefore, we want to find the region *r* that maximizes the likelihood of observing **k**. We wish to compute the contiguous genome region r, where *P* (**K**|*r*) falls above a certain threshold. Assuming Mendelian inheritance, and a Poisson distribution of coverage, for each family, this probability cam be written as:

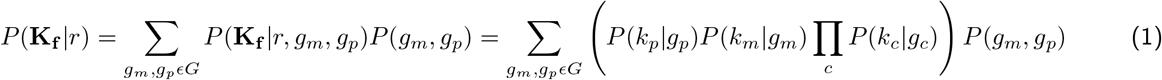

where *k*_*m*_, *k*_*p*_, *k*_*c*_ are the counts of a k-mer in a mother, father, and child. And *g*_*m*_, *g*_*p*_, and *g*_*c*_ are the genotypes of a the mother, father, and child respectively. We can use the phasings to compute the probability of a genotype of a child given their parental genotypes.

We give a full derivation and explanation of assumptions of the maximum likelihood model in 3.

The code for our localization algorithm can be accessed at https://github.com/briannachrisman/alt_haplotypes.

### 5.3 Validation

We validated our algorithm by using datasets of *k*-mer distributions with known locations. We validated two datasets (1) a synthetically generated distribution of *k*-mers and (2) *k*-mer counts extracted from alternative sequences in the decoy reference genome.

We describe how these validation datasets were generated in Supplementary Methods S4. Additionally, our maximum likelihood model is entirely based on biological constants, except for one hyperparameter: *λ*, which determines the length of the region that ASLAN ultimately localizes a subsequence to. We evaluated our algorithm using different values of *λ* on both validation datasets, and found that *λ* = .1 provided a good balance of accuracy (¿90%)and precision (around 1 Mb of resolution). We give a full description of this hyperparameter tuning in S 4.3.

### 5.4 Unmapped Reads Localization and Analysis

#### 5.4.1 Extracting k-mers from Unmapped Reads

We used the previously described pipeline involving jellyfish to extract 100-mers from the unmapped and poorly aligned reads from iHART. To improve computation time, we sampled approximately 100 million out of 200 million *k*-mers (exact number: 104,623,400 out of 234,132,233), under the assumption that many *k*-mers would be overlapping and localize to the same region. We assumed that about 50% sampling would provide sufficient understanding of genetic diversity and missing haplotypes in the T2T assembly, while adhering to computational limitations.

#### 5.4.2 Comparing to Gaps in GRCh38

We retrieved the current gaps in GRCh38 from the GRCh38 AGP file provided by the UCSC genome browser track on Feb 15, 2022. In order to statistically test if our reads localized to gaps more often than by chance, we build a null distribution of the fraction of *k*-mers that localized to gaps in GRCh38 and used it to compute a p-value for the fraction of our *k*-mers that aligned to gaps. To generate the null distribution, for each catalogued gap in the AGP file[11], we randomly generated a new gap on the same chromosome and with the same length as the original gap, but at a different location. We did this for every gap, and then we computed the percentage of unmapped reads that localized to a region containing a gap in GRCh38. We performed this 10,000 times to generate a null distribution for p-value computation.

#### 5.4.3 Comparing to T2T Alignment

We converted the 100-mers from the unmapped reads to fasta format, and aligned them to the T2T’s v1.1 release of T2T-CHM13 (`chm13.draft v1.1.fasta`, accessed on Feb 15, 2022) using bwa-mem with the default parameters. To compare the results of ASLAN with those of alignment to the T2T-CHM13, assembly we translated the starts and ends of our localization coordinates to the T2T-CHM13 coordinate system using `liftOver` [10] and the GRCh38-to-T2T-CHM13 chain file (`grch38.t2t-chm13-v1.1.over.chain.gz`, accessed on Feb 15, 2022)). To prevent split liftOver conversions, we used the `-ends=100000`, for lifting over long sequences.

We considered T2T-CHM13 alignment and ASLAN localization to be in concordance if the liftOver ASLAN localization and the T2T-CHM13 alignment contained an overlapping region.

## 6 Data Access

The raw reads from the iHART samples can be found on Anvil, maintained by NHGRI at https://anvilproject.org/data/studies/ Dataset access is controlled in adherence to NIH Policy and in line with the standards set forth in the individual consents involved in each cohort.

## 7 Acknowledgements

We would like to acknowledge funding from The Hartwell foundation, the Stanford Bio-X Center, the Stanford Precision Health and Integrated Diagnostics Center (PHIND), and the NSF GRFP.

